# The mechanism of Micafungin action to Pteropine orthoreovirus infection in human cell line

**DOI:** 10.1101/2025.01.23.634615

**Authors:** Wirayatida Bubphasook, Atsuo Iida, Eiichi Hondo

## Abstract

Pteropine orthoreovirus (PRV) is a fusogenic virus carried by bats that causes respiratory illnesses in humans. PRVs are transmitted from bats to humans and among humans. Micafungin (MCFG), an approved drug for fungal treatment, shows potential in inhibiting PRV propagation, although its precise mechanism of action remains unclear. This study aimed to investigate molecular mechanisms of MCFG during PRV propagation. Initially, molecular docking was employed to predict the primary target of MCFG. It was the p17 protein of PRV. Prior to examining the effect of MCFG on p17 suppression, differential gene expression analysis was performed to compare PRV-infected host cells with MCFG-treated cells, and it was found that IL-6 should be the main regulator induced by MCFG. Silencing IL-6 with siRNA markedly induced PRV release, syncytial formation, and marginally enhanced PRV RNA replication, corresponding to the main suppressive effects of MCFG against PRV. The p17, the presumed suppression target to inhibit syncytial formation by MCFG, markedly reduced syncytial formation but did not influence viral RNA replication. In contrast, MCFG significantly suppressed syncytial formation and slightly reduced PRV RNA replication, although both MCFG and anti-p17 antibody increased IL-6 mRNA expression. MCFG may inhibit other PRV proteins, like nonstructural replication protein (sigmaNS), which were found by molecular docking study. In conclusion, MCFG primarily targets p17 and modulates host immunity through IL-6, which likely interferes directly with syncytial formation.

## Introduction

Emerging infectious diseases such as severe acute respiratory syndrome coronavirus (SARS-CoV) [1], SARS-CoV-2 [2], Middle East respiratory syndrome coronavirus (MERS-CoV) [3], Ebola virus [4], and Nipah virus [5] are linked to bat-borne pathogens of global significance. These viruses can cause severe illness in humans and result in substantial economic losses across sectors, including healthcare, agriculture, trade, and tourism [6]. Beyond human fatalities, preventing outbreaks and developing effective treatments are critical to reducing the effects of these diseases.

Pteropine orthoreovirus (PRV), belonging to the family *Reoviridae*, is a fusogenic feature and double-stranded RNA virus [7]. The *orthoreovirus* genome is made up of ten double-stranded RNA (dsRNA) segments (L1-L3, M1-M3, and S1-S4). Some fusogenic reoviruses, such as avian orthoreovirus (ARV), broome orthoreovirus (BroV), baboon orthoreovirus (BRV), and PRV, encode two distinctive nonstructural proteins, the fusion-associated small transmembrane (FAST) protein, and p17, located on the S1 or S4 gene segment [8–11]. This virus was first reported in 2006, causing induced respiratory disease in humans, and has been isolated from patients in Malaysia, Indonesia, and the Philippines [12–16]. The PRV exhibits a strong association with the human, leading to the induction of acute respiratory infections (ARI) [17]. Most patients exhibited a complex array of symptoms, including cough, sore throat, and high fever, with some also experiencing diarrhea and vomiting [7]. Until now, the molecular characterizations exhibited that fifteen specific strains of PRV are isolated from *Pteropus poliocephalus*, *P*. *homelands*, *P*. *vampyrus*, *Rousettus leschenaultia*, *R*. *amplexicaudatus*, and *Eonycteris spelaea* in Australia, Malaysia, Indonesia, China, and the Philippines [20–24]. The detection of PRV in fecal samples from monkeys in Thailand indicates its potential role in the transmission of PRV to humans [25]. PRV can also be transmitted from human to human, which could lead to outbreaks and epidemics if not detected and controlled early on [12, 26].

While viral outbreaks pose significant challenges, drug interventions offer promising solutions that can complicate control measures [27]. Drug repurposing seeks to identify new clinical uses for existing drugs with established safety profiles, pharmacokinetics, and other clinical characteristics, reducing the time and costs associated with developing new treatments [27–28]. Micafungin (MCFG), an antifungal drug from the echinocandin class, is derived from fungal secondary metabolites and features a cyclic hexapeptide core with a lipid side chain [29]. MCFG specifically targets the fungal cell wall by disrupting proteins responsible for synthesizing β-1,3 glucan polysaccharides [30]. Approved by the United States Food and Drug Administration (US FDA) for treating *Candida sp*. infections [31], MCFG has also shown antiviral activity against enterovirus 71 [32], chikungunya virus [33], dengue virus [34], and SARS-CoV-2 [35].

Previous work reported that MCFG exhibits antiviral activity against PRV infection in human cell lines by effectively inhibiting viral release and syncytial formation [36]. However, the evident mechanism for micafungin inhibiting PRV remains unknown. This study aims to clarify the molecular mechanism of MCFG in PRV propagation. The molecular docking was conducted to predict the target of MCFG against the PRV protein. Following this, gene expression profiles were analyzed to study the effect of the target PRV protein on host cell responses by next-generation sequencing. A candidate gene was knocked down by siRNA to investigate its role in PRV replication. Lastly, an antibody against the target protein of MCFG was generated to explore the effect of MCFG through viral inhibition assays. Understanding the molecular mechanisms of MCFG is crucial for advancing antiviral therapy, ensuring the development of effective treatments, and preventing future outbreaks.

## Materials and methods

### Molecular docking

The PRV50G protein sequences were obtained from the NCBI database (https://www.ncbi.nlm.nih.gov) as shown in Supplementary Information 1A, and the viral protein structure was modeled using Alphafold2 (https://alphafold.ebi.ac.uk) [37]. The 2D structure of micafungin was sourced from the PubChem database (https://pubchem.ncbi.nlm.nih.gov), and its 3D structure was generated and optimized for geometry using Avogadro (v1.2.0) [38]. Molecular docking experiments were conducted to target the PRV proteins using AutoDock 4.2.6 (http://autodock.scripps.edu/) [39–41]. During the docking process, explicit hydrogens, charges, and flexible torsions were assigned to the protein and the ligands. Polar hydrogens were added to ensure the correct ionization and tautomeric states of the amino acid residues of the protein, and Kollman united atom charges were applied to the protein. The rigid roots of ligands were identified, with rotatable bonds set to allow flexibility. The modified 3D structures of the PRV protein and ligands, which incorporated their bond flexibility, were converted to the PDBQT format required for AutoDock calculations. The Lamarckian genetic algorithm (LGA) was employed to explore potential binding conformations using the following settings: 300 dockings in the population, a maximum of 27,000 generations, 25 million energy evaluations, and 50 docking runs with randomized initial positions and conformations. Default parameters were applied for other settings, including mutation and crossover rates. AutoGrid was used to define the search space by generating a grid encompassing the entire protein structure. Pre-calculated grid maps representing interaction energies between ligand atom probes and the receptor were generated with AutoGrid 4.2. The conformation with the lowest binding energy was selected as the most favorable binding pose. All docking results, including images and outputs from AutoDock, were further analyzed using UCSF Chimera (https://www.cgl.ucsf.edu/chimera/) [42].

### Cells

Vero JCRB9013 (African green monkey kidney cells) and A549 (Human lung tissue cells) were used in this study. Both cell lines were cultured and maintained in Dulbecco’s Modified Eagle’s Medium (DMEM; Nissui, Tokyo, Japan) containing 10% heat-inactivated fetal bovine serum (FBS; HyClone, Logan, UT), 2 mM L-glutamine (Sigma-Aldrich, St. Louis, MO), 0.14% sodium bicarbonate (NaHCO□; Sigma-Aldrich, St. Louis, MO), and 100 U/mL-0.1 μg/mL penicillin-streptomycin (Meiji, Tokyo, Japan) at 37°C in a humidified atmosphere with 5% CO□.

#### Pteropine orthoreovirus

The *Pteropine orthoreovirus* strain Garut-50 (PRV50G), previously isolated from the greater flying fox (*Pteropus vampyrus*) in Indonesia [24], was propagated in Vero cells in DMEM medium containing 2% FBS and stored at −80°C until use. Virus titration was performed by plaque assay in the Vero 9013 cell line.

### Micafungin

Micafungin sodium for intravenous infusion (Nipro Corporation, Osaka, Japan) was dissolved in sterile DMSO at 10 mM to prepare stock solutions and stored at −30°C until use.

### Virus infection, drug adding, and RNA extraction

A549 cells were seeded onto a 6-well plate at a density of 1×10□ cells per well and incubated in a DMEM medium containing 10% FBS for 24 hours to form a monolayer. The medium was removed the following day, and the cells were washed twice with 1X Phosphate-Buffered Saline (PBS). Then, a DMEM medium containing 2% FBS was added to the cells, and they were infected with PRV50G at a multiplicity of infection (MOI) of 0.1. At 2 hours post-infection (hpi), the PRV was removed, and micafungin was added to the cells at a final concentration of 20 µM. DMEM medium containing 2% FBS was used as a control. All treatments were performed in triplicate. After 6 hours of infection, RNA was extracted using ISOGEN2 (Nippon Gene, Tokyo, Japan). Total RNA was stored at - 80°C until used for RNA sequencing.

### RNA-Seq

RNA samples were sent for next-generation sequencing, which was outsourced to Macrogen (Kyoto, Japan). The analysis of the FASTQ files was performed using the supercomputer system at the National Institute of Genetics (Shizuoka, Japan). First, quality checks and adapter trimming (using the -g option) were conducted with Fastp software (v0.20.1). For mapping, the human reference genome GRCh38.p13 (National Center for Biotechnology Information; NCBI) was used, and gene annotations were obtained from GENCODE v42 annotation [43]. The human reference sequence, Homo_sapiens.GRCh38.cdna.all.fa was downloaded from Ensembl. Mapping was performed using HISAT2 software (v2.1.0, with default settings), followed by conversion of the SAM files to BAM format using samtools (v1.7) [44]. Read counts were obtained for humans using featureCounts (v2.0.1). Differentially expressed genes were identified using the DESeq2 package (v1.36.0) in R (v4.3.2) [45]. Gene ontology (GO) analysis of significantly upregulated genes was conducted using the clusterProfiler package (v4.10.0) [46]. The Venn diagram was created using the web-based tool Venny 2.1 (https://bioinfogp.cnb.csic.es/tools/venny/). Protein-protein interactions were analyzed using the STRING database (v12) (https://string-db.org) [47].

### RT-qPCR

All RNA samples were adjusted to a final 50 ng/µl concentration. A 4 µl aliquot of each RNA sample was mixed with 2 µl of nuclease-free water. The solutions were incubated at 65°C for 5 minutes and then placed on ice for at least 5 minutes. A mixture containing 2 µl of 5X RT buffer, 0.5 µl of 10 mM dNTPs, 0.5 µl of 10 µM Oligo(dT)20, and 0.5 µl of ReverTra Ace™ (TOYOBO, Osaka, Japan) was added to the RNA solutions. The RNA solutions were then incubated at 42°C for 60 minutes, followed by 99°C for 5 minutes. The cDNA was stored at - 80°C until required for further analysis. The expression of candidate genes was determined by real-time qPCR using specific primers, as shown in Supplementary Information 1B. THUNDERBIRD™ Probe and SYBR® (TOYOBO, Osaka, Japan) qPCR Mix was used as the reaction master mix, optimized for dye-based quantitative PCR (qPCR). For cDNA amplification, 1 µl of cDNA was combined with 5 µl of THUNDERBIRD™ Probe qPCR Mix, 0.5 µl of 10 µM specific gene forward and reverse primers, and 3 µl of nuclease-free water, adjusted to a final volume of 10 µl. The thermal cycling was performed using a LightCycler® (Basel, Switzerland). Gene expression was analyzed using the 2^-ΔΔCt^ comparative method [48] to determine fold differences.

### siRNA and siRNA transfection

The sequences of the siRNAs are as follows: siRNA IL-6 sense: 5’-GGACAUGACAACUCAUCUCtt-3’, siRNA IL-6 antisense: 5’-GAGAUGAGUUGUCAUGUCCtt-3’, siRNA scrambled control sense: 5’-AUUGGGUAGUGUUUCAGGCtt-3’, and siRNA scrambled control antisense: 5’-ACGUGACACGUUCGGAGAAtt-3’ (FASMAC, Kanagawa, Japan). Polyethylenimine (PEI) was used as the transfection reagent for knockdown in the cell line, following the protocol by Sabrina Höbel and Achim Aigner (2013) [49]. A total of 250 pmol of siRNAs were transfected into A549 cells for 8 hours, after which the siRNAs were removed. Only the transfection reagent was used as the control. A DMEM containing 2% FBS was added to the transfected cells and incubated for 24 hours post-transfection. Next, PRVs were added to the medium. At 6 hpi, four randomly selected fields were examined under a light microscope, and syncytia areas were quantified using ImageJ. Total RNA was extracted from cell pellets and stored at −80°C until used for gene expression and PRV copy number analysis. Supernatants were also stored at −80°C until TCID_50_ measured viral titers. All treatments were performed in triplicate.

### Virus detection

Total RNA from infected cells was extracted using ISOGEN2. RNA was reverse-transcribed into cDNA and subjected to qPCR using RNA-direct SYBR Green Realtime PCR Master Mix (TOYOBO, Osaka, Japan) under the following conditions: 90°C for 30 seconds, 61°C for 20 minutes, 95°C for 30 seconds, followed by 35 cycles of 95°C for 5 seconds, 55°C for 10 seconds, and 74°C for 15 seconds. The melting curve analysis was conducted with steps at 95°C for 10 seconds, 65°C for 60 seconds, and 97°C for 1 second. Primer sequences were designed to amplify a region of the S4 segment of PRV50G (forward: 5′-TTGGATCGAATGGTGCTGCT-3′; reverse: 5′-TCGGGAGCAACACCTTTCTC-3′; amplicon size: 159 bp). A standard curve was generated, and Ct values were plotted. The relative viral RNA copy number compared to the control was determined by dividing the values of the experimental group or control (vehicle) by the average value of the vehicle group. These results were then logarithmically transformed, and the mean and standard deviation were calculated.

### TCID_50_

The Vero JCRB9013 cells were seeded in a 96-well cell culture plate at a density of 5×10− cells per well and incubated for 24 hours to form a monolayer. The following day, a vial of frozen PRV was rapidly thawed at 37 °C for 90 seconds, and the virus was 10-fold serially diluted from 10□¹ to 10□¹□ in DMEM containing 2% FBS. After dilution, 100 µl of each viral dilution was incubated on the Vero JCRB9013 cells with eight replicates. The last two columns of the cell line were used as negative controls. The cytopathogenic effect (CPE) was observed daily at each dilution. After 1 week of infection, each well was fixed with 50 µl of 4% formalin (v/v) in 1X PBS. The solution was then discarded, after which 1% crystal violet (w/v) in 1X PBS was added to each well to stain the live cells. Afterward, the number of CPEs was counted, and the TCID_50_ was calculated using the Spearman-Kärber method.

### Quantification of syncytial formation

At 6 hpi, four randomly selected fields were captured using a light microscope at 200x total magnification. All treatments were performed in triplicate. Syncytia formation in the infected cells was quantified using ImageJ software. The thresholds were adjusted by covering the syncytial areas and measuring them to determine their total surface area. Then, the image was converted to a binary format to separate adjacent syncytia that might appear as a single mass. The images were analyzed by particles by using the default setting. The measured values were then normalized to the corresponding mock-treated controls to account for background variations. Results were presented as the relative syncytial area normalized to the control condition.

### Recombinant antibodies against PRV p17 protein

The pET-46 Ek_LIC vector was used for construction. The BseRI site was used for insertion. A full length of p17 was generated by RT-PCR. First, the PRV RNA was extracted and converted to cDNA. Then, cDNA was amplified with primer p17_BseRI forward; 5’ ACGTGGATGACGACGACAAGATGTCCATCCAGCCTCATCT 3’ and p17_BseRI reverse; 5’ ATTTGAGGAGAAGCCCGGACTCAGATCGCGAAGCGCTTAT 3’. After amplification, the entire length of p17 was inserted into the vector using the NEBuilder® HiFi DNA Assembly Cloning Kit (New England Biolabs, Ipswich, MA). The NEB 5-alpha chemically competent cells were used for cloning. Then, plasmids were extracted using the FavorPrep Plasmid Extraction Mini Kit (FAVORGEN, Taiwan). Next, plasmids were transformed into *Escherichia coli* BL21 (DE3) (Thermo Fisher Scientific, Waltham, MA) for the expression of recombinant proteins. The recombinant clones were cultured on Luria-Bertani (LB) agar containing 50 µg/mL ampicillin at 37 °C. The single recombinant clone was later inoculated in LB broth with the same antibiotics and cultured until OD600 reached 0.4-0.6. Isopropyl-β-D-thiogalactopyranoside (IPTG) was added into a bacterial culture to the final concentration of 0.3 mM, and the culture was incubated at 37°C for 3 hours with vigorous shaking. After induction, the recombinant cells were pelleted by centrifugation. The cell pellet was resuspended with 30 µl of 1X PBS, 25 µl of 2X SDS-PAGE sample buffer, and 5 µl of β-mercaptoethanol. Next, the cell suspension was boiled for 10 min. After boiling, the sample was centrifuged at 13,000 rpm at 4 °C for 10 min. The supernatant was loaded in 15% SDS-PAGE. A gel was stained by Coomassie Brilliant Blue staining solution and de-stained in nuclease-free water. For protein purification, proteins were extracted from the cell pellet by sonication on ice for 2 minutes in binding buffer (20 mM sodium phosphate, 0.5 M sodium chloride, 20 mM imidazole, pH 7.4), then centrifuged at 7,000 rpm for 20 min. The supernatant was collected and filtered through a 0.45-micron polyvinylidene difluoride (PVDF) sterilization membrane syringe filter before protein purification by Cytiva™ HiTrap™ Chromatography Columns (Marlborough, MA). The column was washed with a binding buffer, and the sample was eluted with an elution buffer (binding buffer containing 500 mM imidazole). The elution buffer was later changed to a 1X PBS using an Amicon® Ultra-15 Centrifugal Filter Unit (Merck Millipore, Darmstadt, Germany). Fractions containing purified protein were pooled and stored at 4 °C. The affinity purified recombinant p17 protein of 50 µg was mixed with TiterMax® (TiterMax USA Inc., Norcross, GA) Gold Adjuvant in a 1:1 ratio to a final volume of 100 µL for 15 minutes at room temperature. Then, the mixture was injected subcutaneously under the back skin of the ICR mice to stimulate an immune response (the Ethics Review Board for Animal Experiments was approved at Nagoya University, approval number A240118-004). After two weeks (Day 14), mice received a booster dose consisting of the same protein mixed with an adjuvant. The booster was administered in the same manner as the primary injection. A final booster was administered on day 42 (six weeks after the initial immunization). This final injection consisted of the recombinant p17 protein without any adjuvant to avoid prolonged immune stimulation. This boost was given 3-4 days before blood collection. Periodic blood collections were conducted once a week before the final dose. On days 45-46, blood samples were collected from each mouse by cardiac puncture under deep anesthesia to ensure minimal pain and discomfort. The titers of serum were determined by the Enzyme Linked Immunosorbent Assay (ELISA) method. The sera of immunized mice were used for functional viral inhibition assays.

### ELISA

2.5 µg of p17 recombinant protein were coated onto microplates for ELISA and incubated overnight at 4 °C. The next day, the coated wells were washed three times with 100 µl of Phosphate Buffered Saline-Tween 20 (PBS-T). Then, 5% skim milk was added to block non-specific binding for 1 hour. Serum from immunized mice, diluted 1:10, 1:100, 1:500, and 1:1000, was added to the wells and incubated for 2 hours. Serum from non-immunized mice was used as a control. After incubation, the wells were washed three times with PBS-T, and Horse Anti-Mouse IgG Antibody (H+L), Peroxidase (Vector Laboratories, Newark, CA), diluted 1:1000, was added to each well and incubated for 1 hour. Afterward, the wells were washed five times with PBS-T. Finally, 100 µl of ABTS® 2-Component Microwell Peroxidase Substrate was added to each well and incubated for 10 minutes. The absorbance at 405 nm was then read using an iMark™ Microplate Absorbance Reader (Bio-Rad Laboratories, Inc., Hercules, CA) to measure the signal of the p17 protein.

### Functional viral inhibition assay

A549 cells were seeded into a 12-well plate at a density of 1 × 10□ cells per well to form a monolayer. The next day, the medium was removed, and the cells were washed twice with 1X PBS. They were then infected with PRV50G at a MOI of 0.01 in DMEM containing 2% FBS. At 2 hpi, the PRV was removed, and anti-serum, MCFG, or a mixture of anti-serum and MCFG was added to the cells. Mock and PRV-infected cells were used as a control. All treatments were performed in triplicate. Six hours post-infection, random areas of cells were imaged under a microscope, and syncytia areas were quantified using ImageJ. Total RNA was extracted from cell pellets and stored at −80°C until it was used to determine PRV copy number and IL-6 gene expression.

### Statistical analysis

All graphs in this study were created, and the statistical analysis of the relative mRNA expression levels and virus copy number was conducted using a non-parametric t-test in Prism 8 software (GraphPad Software Inc., San Diego, CA). A *p*-value of less than 0.05 was considered as significantly different, and *p*-values were indicated with n.s.; not significant, *; *p* < 0.05, **; *p* < 0.01, ***; *p* < 0.001, **** ; *p* < 0.0001.

## Results

### p17 is a target of MCFG

To predict the potential target of micafungin, the *in silico* docking was analyzed, revealing the p17 non-structural protein (p17 nsp) exhibited the highest binding efficiency among the 12 tested PRV proteins. The p17 demonstrated the most favorable binding conformation with MCFG, attaining a binding energy of −4.23 kcal/mol, succeeded by the nonstructural replication protein (sigmaNS gene, −2.35 kcal/mol), major-inner capsid protein (sigma1 gene, −1.32 kcal/mol), the major-inner capsid protein (lambdaA gene, −1.17 kcal/mol), and the membrane fusion protein (−0.07 kcal/mol), as depicted in Fig. 1A. The binding site of micafungin with p17 is depicted in Fig. 1B. The three-dimensional and two-dimensional configurations of amino acid residues engaged in the interaction with the MCFG ligand demonstrated conventional hydrogen bond interactions with HIS49, ARG90, ILE70, ARG71, THR116, LYS122, HIS125, and CYS126, as well as carbon-hydrogen bond interactions with PHE127 (to the right of Fig 1B).

**Fig. 1.**
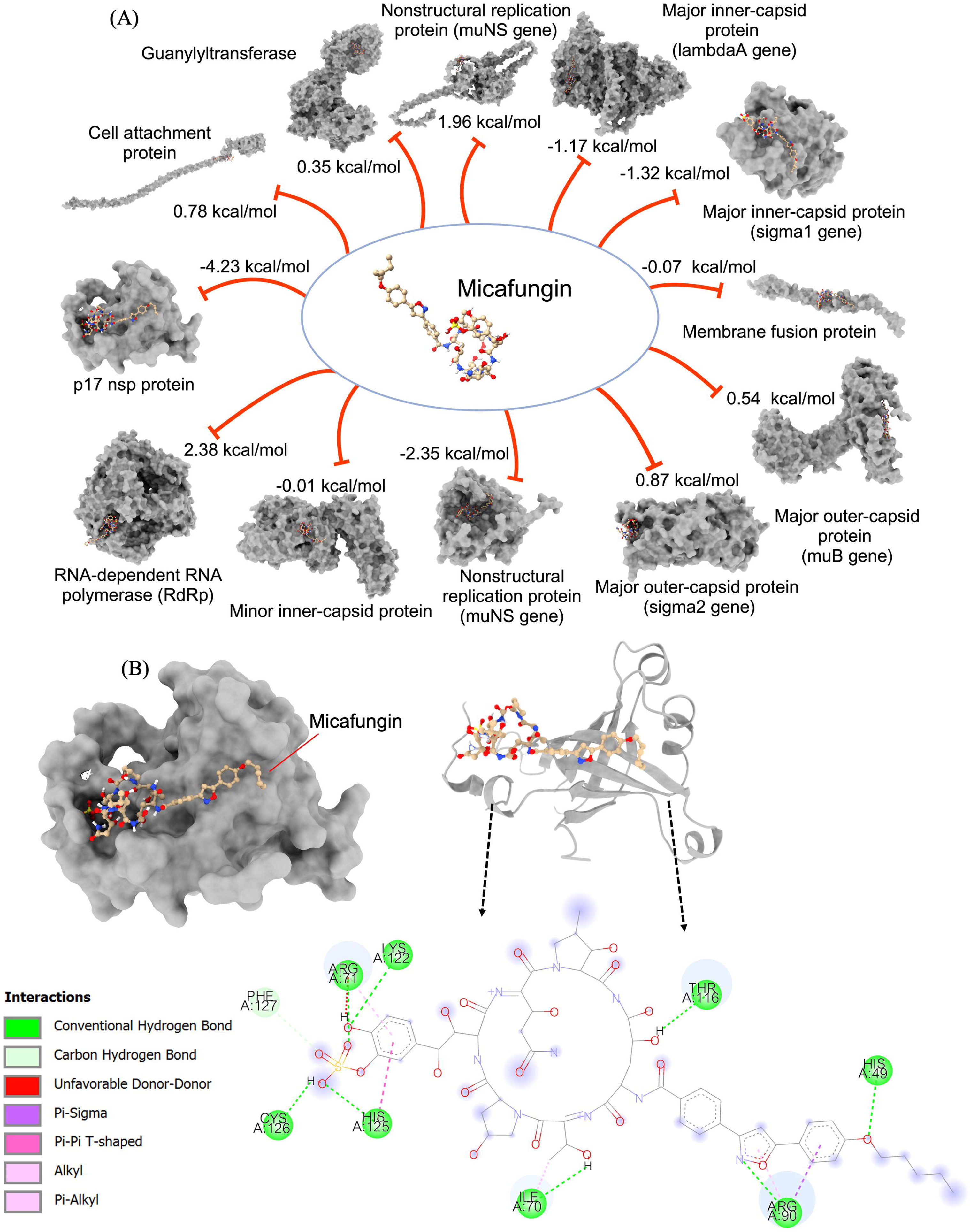
Illustration of micafungin (MCFG) inhibition against nine PRV target proteins with binding energy results from AutoDock (A). Docking interactions of MCFG with the p17 nsp are shown, including the surface structure of the best binding mode within the protein pocket (left), the 3D structure of amino acid residues involved in the interaction with the MCFG ligand (top-right), and the 2D representation of MCFG binding interactions (down-right), highlighting conventional hydrogen bonds (Dark green dashed lines), Carbon-hydrogen bonds (Light green dashed lines) (B).

### The syncytial formation

The syncytial formation is one of the morphological markers to evaluate the PRV pathogenic and micafungin effect. The time course of syncytial formation following PRV infection in A549 cells was examined to identify the molecular pathways related to the micafungin administration. Syncytia began forming at approximately 6 hpi in PRV treatment. Additionally, MCFG exhibited syncytial formation at 6 hpi (Fig. 2A).

**Fig. 2.**
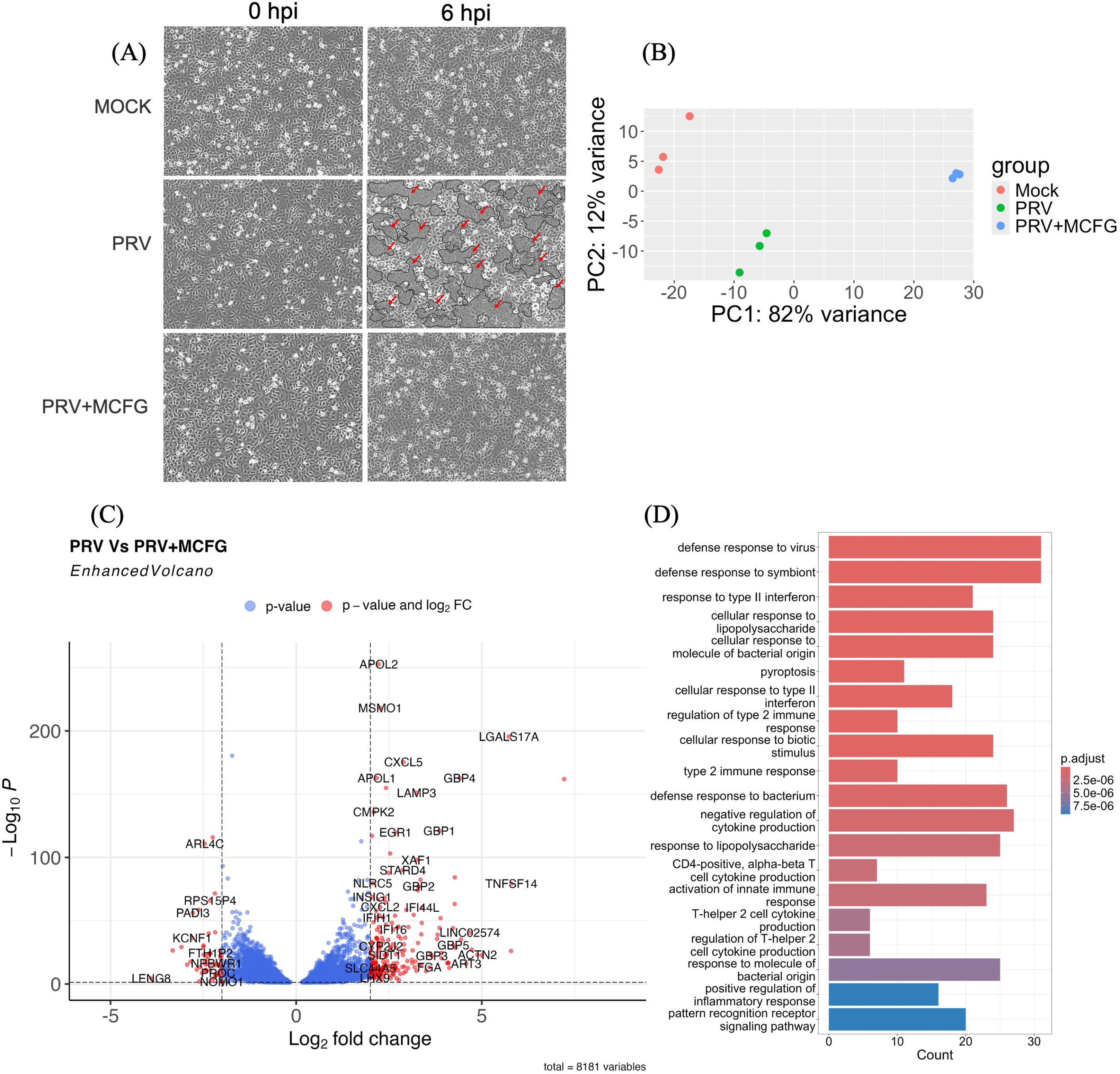
Morphology of A549 infected with PRV and PRV treated with 20 μM of MCFG at 6 hpi. The marked area with arrows indicates syncytial formation (A). The PCA analysis of RNA-seq data was used to visualize sample to sample variation with 3 replicates (B). Volcano depicting differentially expressed genes in the A549 cell line infected with PRV and PRV treated with MCFG while red dots represent genes with -log_10_P values more than 0.05 and log2 fold change values more than 2 (C). Gene Ontology (GO) enrichment of DEGs between PRV and PRV treated with MCFG (D).

### Gene profiles by next-generation sequencing

The principal component analysis (PCA) was conducted among the mock, PRV-infected, and PRV-treated with MCFG groups (Fig. 2B). Each independent replicate clustered closely within its respective treatment group; however, the mock and micafungin-treated samples exhibited greater dispersion than the PRV-infected samples. Principal Component 1 (PC1) represented 82% of the variance, distinguishing the mock and PRV-infected samples from the micafungin-treated group, while Principal Component 2 (PC2) showed 12% of the variance between the mock and micafungin-treated samples and those affected by PRV infection. A total of 203 differentially expressed genes (DEGs) were identified between PRV-infected samples and those treated with MCFG, with the indoleamine 2,3-dioxygenase 1 (IDO1) gene exhibiting the most significant upregulation. Conversely, 58 genes were downregulated, with the leukocyte receptor cluster member 8 (LENG8) gene exhibiting the lowest expression level (Supplementary Information 1C). These DEGs are displayed in a volcano plot (Fig. 2C). Gene ontology (GO) enrichment of DEGs was performed for further functional analysis, focusing on the biological process (BP) category. The top 20 enriched BP terms are presented in Fig. 2D, with the most significantly enriched terms being “defense response to viruses” and “defense response to the symbionts”.

A comparative analysis of DEGs was performed to identify genes associated explicitly with MCFG treatment during PRV infection. Venn diagram analysis compared DEGs between mock-treated and PRV-infected cells (MOCK vs. PRV) and between PRV-infected cells and PRV-infected cells treated with MCFG (PRV vs. PRV + MCFG), highlighting genes uniquely expressed in MCFG-treated PRV-infected cells. For the upregulated gene, 147 unique genes were identified (Fig. 3A), with their details provided in Supplementary Information 1D. GO enrichment analysis revealed that the top enriched biological processes among the upregulated genes were “response to lipopolysaccharide” and “defense response to bacterium” (Fig. 3B). Protein-protein interaction (PPI) network analysis (Fig. 3C) identified interleukin 6 (IL-6) gene as a central hub, exhibiting the highest number of interactions and being closely associated with enriched GO terms (Fig. 3D).

**Fig. 3.**
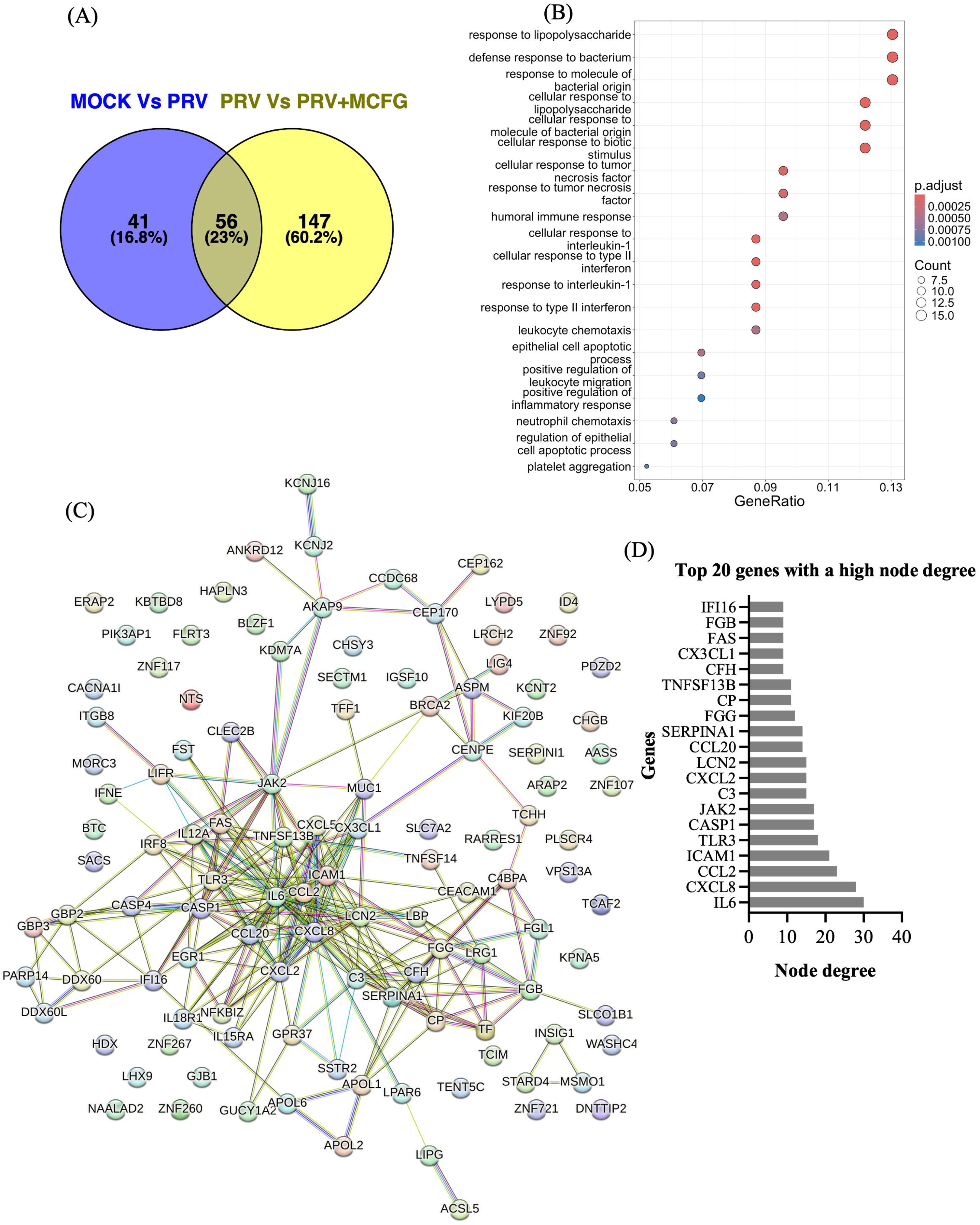
A Venn diagram showing the differentially upregulated genes between Mock and PRV, compared to PRV and PRV treated with MCFG (A). The top 20 enriched biological process pathways with a *p*-value < 0.001 are labeled on the y-axis. The x-axis represents the Z-score values, indicating the degree of enrichment (B). PPIs of the DEGs by the STRING database, representing interaction pathways derived from 147 genes identified across all selected gene lists (C). A bar graph depicting the top 20 genes with the highest node degree values from the STRING protein interaction network, highlighting their central role in the interaction network (D).

A total of 58 unique downregulated genes were identified (Fig. 4A), with further details in Supplementary Information 1D. GO enrichment analysis highlighted “intramembranous ossification” and “direct ossification” as the most enriched biological processes (Fig. 4B). PPI analysis revealed the ribosomal protein L9 (RPL9) gene as the central node among the downregulated genes, exhibiting the highest degree of interactions (Fig. 4C). The top 10 downregulated genes with the highest node degree are presented in Fig. 4D.

**Fig. 4.**
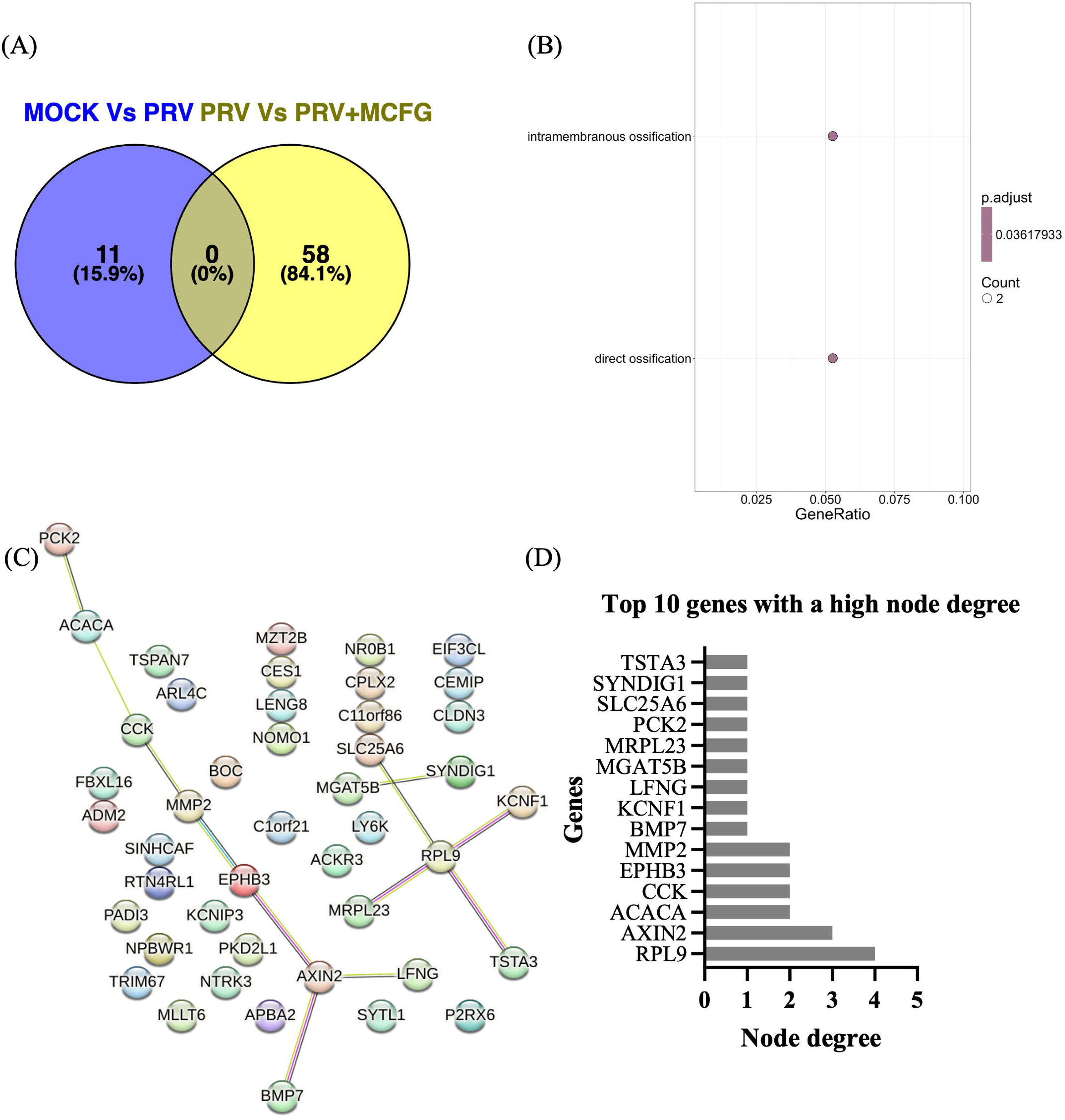
A Venn diagram showing the differentially downregulated genes between Mock and PRV, compared to PRV and PRV treated with MCFG (A). The top enriched biological process pathways with a *p*-value< 0.001 are labeled on the y-axis. The x-axis represents the Z-score values, indicating the degree of enrichment (B). PPIs of the DEGs by the STRING database, representing interaction pathways derived from 58 genes identified across all selected gene lists (C). A bar graph depicting the top 20 genes with the highest node degree values from the STRING protein interaction network, highlighting their central role in the interaction network (D).

Among the top 20 upregulated genes in MCFG-treated PRV-infected cells, IL-6 was chosen for further examination because of its significant role in immune response and its pivotal position in the PPI network. Further research concentrated on clarifying the function of IL-6 in the mechanism of PRV propagation and its modulation by MCFG.

### IL-6 knockdown

The MCFG showed it can induce IL-6 gene expression when compared to PRV infection (*p* < 0.05; Fig. 5A), and it also slightly inhibits PRV RNA replication (*p* < 0.01; Fig. 5B) while considerably decreasing PRV release (*p* < 0.05; Fig. 5C) and syncytial formation (*p* < 0.0001; Fig. 5D). To investigate the function of IL-6 in PRV propagation, IL-6 gene knockdown was performed utilizing siRNA directed against IL-6 prior to PRV infection. Knockdown efficiency analysis revealed significant downregulation of IL-6 expression in siRNA-treated cells compared to PRV-infected cells without siRNA treatment (*p* < 0.01) and those treated with scrambled siRNA control following PRV infection (*p* < 0.0001; Fig. 5A). Viral copy numbers, measured by RT-qPCR, were markedly elevated in IL-6 knockdown cells compared to PRV-infected cells lacking siRNA (*p* < 0.05) or those treated with scrambled siRNA control (*p* < 0.05; Fig. 5B). Similarly, the virus titer in the supernatant, measured using the TCID□□ assay, was significantly elevated in IL-6 knockdown cells compared to PRV-infected cells (*p* < 0.01) and scrambled siRNA controls (*p* < 0.01; Fig. 5C). In addition, IL-6 knockdown substantially enhanced syncytial formation. Quantitative analysis showed a significant increase in syncytial formation in IL-6 knockdown cells compared to PRV-infected cells (*p* < 0.0001) and scrambled siRNA controls (*p* < 0.0001; Fig. 5D). The IL-6 gene was identified as highly interconnected with other genes within the PPI network (Fig. 2C). Thus, the effect of IL-6 knockdown on the expression of related genes was examined. Tumor necrosis factor superfamily member 13B (TNFSF13B) and intercellular adhesion molecule 1 (ICAM1) were slightly downregulated in IL-6 knockdown cells compared to scrambled siRNA-treated cells *(p* < 0.05; Figs. 5E and 5F).

**Fig. 5.**
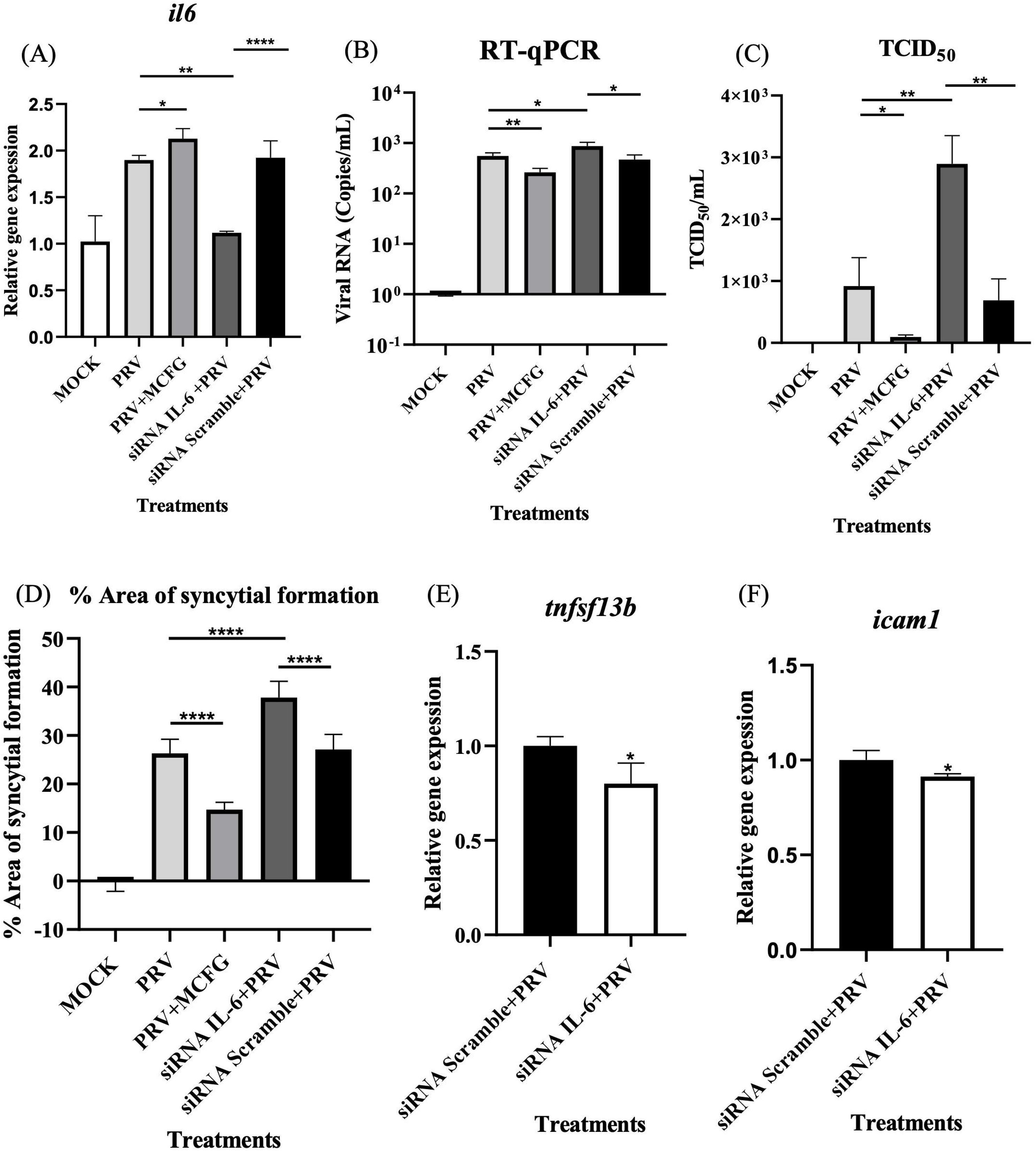
IL-6 gene expression analysis with siRNA against IL-6 and MCFG. Values were calculated using the 2^-ΔΔCt^ comparative method. Statistical differences between groups by a student’s t-test. Bars represent the mean (n=3) relative gene expression from three independent experiments with standard deviation (*; *p*-value < 0.05, **; *p* < 0.01, ***; *p* < 0.001, ****; *p* < 0.0001) (A). Viral copy numbers by qPCR, with data represented as the mean (n=3) (B). Virus titer in the supernatant by TCID□□ assay (n=3) (C). The percentage of syncytial formation area was calculated using ImageJ software (n=12) (D). TNFSF13B gene expression analysis with siRNA against IL-6 and MCFG (E), and ICAM1 gene (F).

### Inhibition of p17 protein

Anti-p17 antibody in PRV-infected cells notably reduced the area of syncytial formation when compared to cells infected with either normal sera (*p* < 0.0001) or PRV alone (*p* < 0.001). The combination of the MCFG drug and the anti-p17 antibody led to a significant decrease in syncytial formation compared to either treatment alone, with *p* < 0.0001 for the drug alone and *p* < 0.0001 for the anti-p17 antibody alone (Fig. 6A). The anti-p17 antibody did not substantially influence viral RNA replication when compared to normal sera or PRV alone, as illustrated in Fig. 6B. Likewise, both MCFG and anti-p17 sera exhibited substantial decreases in syncytial formation in a dose-dependent manner, with 10 µM of MCFG and anti-p17 antibody at titer 10 being the most highly inhibited syncytial formation (Fig. 6C). The expression of the IL-6 gene was significantly upregulated in samples treated with anti-p17 antibody compared to those treated with PRV alone (*p* < 0.001) and the normal serum control (*p* < 0.001), as depicted in Fig. 6D.

**Fig. 6.**
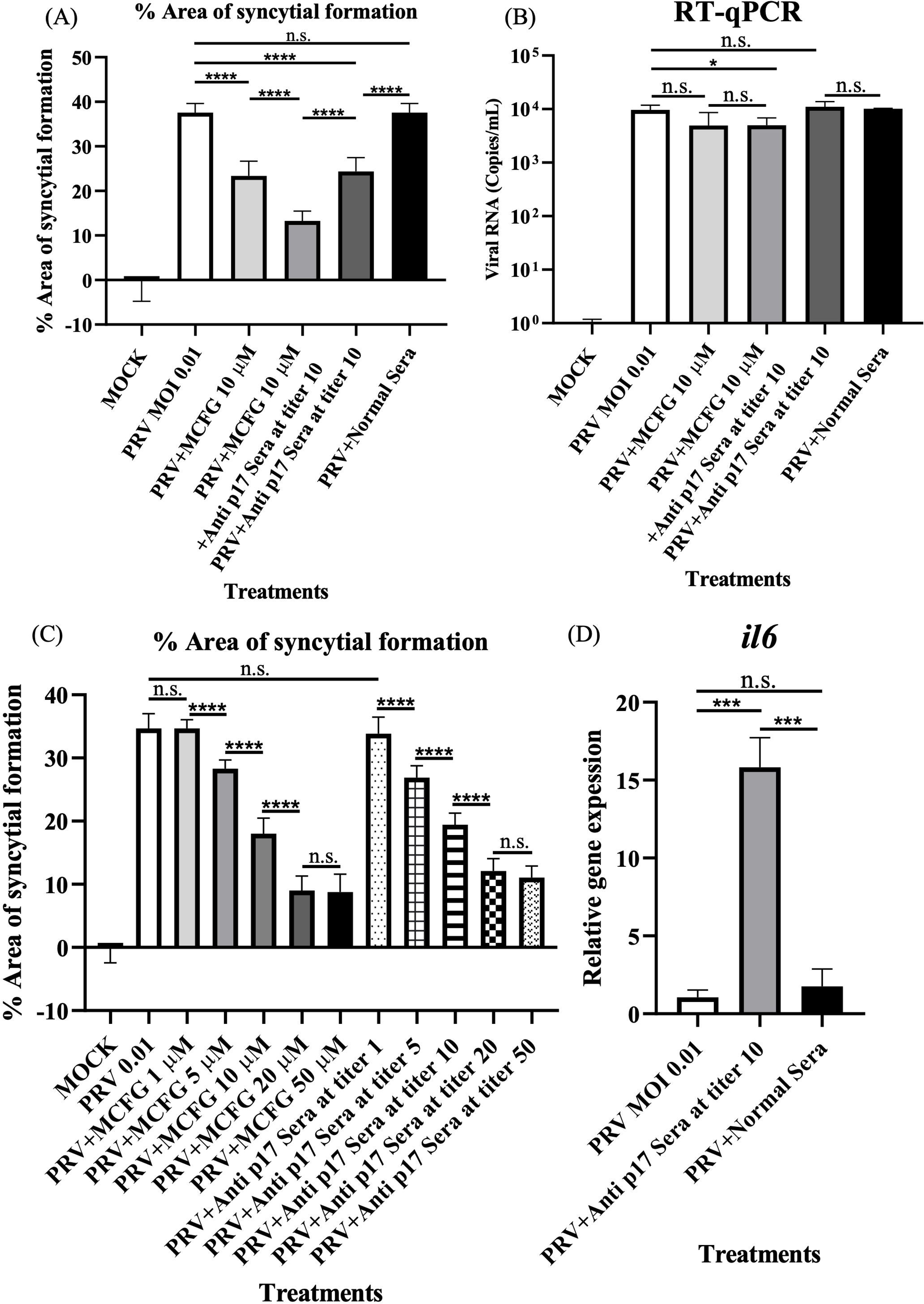
Percentage of syncytial formation area by ImageJ software (n=12). Statistical differences between by a student’s t-test. Bars represent the mean (n=3) relative gene expression from three independent experiments with standard deviation (*; *p*-value < 0.05, **; *p* < 0.01, ***; *p* < 0.001, ****; *p* < 0.0001) (A). Viral copy numbers by qPCR (n=3) (B). Percentage of syncytial formation area across different concentrations of MCFG and anti-p17 sera (n=12) (C). IL-6 gene expression (n=3) (D).

## Discussion

Micafungin is recognized for its ability to inhibit 1,3-β-glucan synthase, an enzyme that plays a crucial role in fungal cell wall synthesis [30]. Besides this function, MCFG and its analogs, including caspofungin and anidulafungin, have established antiviral properties against several viruses, such as enterovirus 71, chikungunya virus, dengue virus, Zika virus, SARS-CoV-2, and PRV [32–34, 50, 51, 36]. This study investigated the mechanism of MCFG during PRV propagation in human cell lines.

The IL-6 gene, a pleiotropic cytokine that has long been recognized for its central role in immune modulation and response to infections, was selected as the candidate gene in this study. IL-6 is produced in response to tissue damage and infections. It exerts its effects through the JAK/STAT3 signaling pathway, which influences various immune responses, including inflammation, T-cell differentiation, and tissue repair [52]. IL-6 is particularly important in coordinating pro-inflammatory and anti-inflammatory responses, making it a critical player in controlling viral replication and immune function [53, 54]. In the context of viral infections, IL-6 helps regulate the immune system by enhancing cytotoxic T-cell activity and IgG production and by promoting tissue repair while reducing viral-induced apoptosis [55]. IL-6 has been shown to play an essential role in limiting viral replication in several other viral infections, such as hepatitis B virus (HBV), by inhibiting viral replication and interfering with viral receptor expression [56, 57]. PRV7S-infected NP460 cells reduced IL-6 cytokine expression. The precise mechanism underlying the downregulation of IL-6 in nasopharyngeal cells following PRV7S infection remains inadequately elucidated [58]. Our study demonstrates that IL-6 plays a pivotal role in slightly suppressing PRV RNA replication, significantly reducing viral release and syncytial formation in the A549 cell line, the same as the effect of MCFG against PRV (Fig. 5), consistent with the primary suppressive effects of MCFG on PRV. The suppression of the IL-6 gene significantly led to the upregulation of TNFSF13B and ICAM1 (Figs. 5E, 5F). These changes, which are linked to IL-6, suggest that IL-6 should act as a central regulator of the host response to micafungin treatment during PRV propagation. This implies that IL-6 modulates a network of genes involved in antiviral defense and cellular responses, underscoring its critical role as a key mediator in inhibiting PRV propagation by MCFG.

The nonstructural orthologous proteins ARV p17, BRV p16, BroV p16, and PRV p17 are encoded by the second open reading frame (ORF) of the fusogenic reovirus S1 or S4 gene segment. [8–11, 59]. Previous findings indicate that the ARV p17 functions as a CRM1-independent nucleocytoplasmic shuttling protein, possessing a nuclear localization signal defined by essential basic amino acids, K122 and R123, which are conserved among ARV [60]. The ARV p17 has demonstrated the ability to activate the p53 signaling pathway while inhibiting the PI3K/AKT/mTOR and ERK pathways. These interactions lead to impaired cellular translation, cell cycle arrest, and autophagosome formation, facilitating effective viral replication [61–65]. Specifically, PRV p17 plays an important role in enhancing FAST-mediated cell-cell fusion or syncytial formation, a key process for viral spread in bat cells. However, overexpression of PRV p17 can lead to excessive fusion, resulting in host cell damage. This observation points to the presence of a regulatory mechanism that controls PRV p17 expression, possibly through interactions with host translation machinery, which may be bat-specific [66]. Our study highlights that MCFG was predicted to bind to the p17 nsp of PRV and that the addition of anti-p17 antisera at titer 10 significantly reduced syncytial formation compared to PRV infection alone (Fig. 7A), while viral RNA replication remained unaffected (Fig. 7B). This is consistent with previous reports suggesting that p17 is not essential for PRV replication in A549 cells [66]. The combination of 10 µM MCFG and anti-p17 sera at a titer of 10 significantly enhanced the inhibition of syncytial formation compared to either treatment alone (Fig. 7A). The enhanced effect suggested that MCFG and anti-p17 antibodies synergistically inhibited the p17 protein, which promotes cell-cell fusion. Syncytial formation was dose-dependently reduced with higher concentrations of MCFG or anti-p17 sera. The strongest effect was observed with 20 µM MCFG and anti-p17 sera at a titer of 20 (Fig. 7C). Consequently, specific antibodies targeting p17 increased IL-6 expression in host cells compared to those treated with PRV alone or PRV with normal sera (Fig. 7D). These indicate that the suppression of p17 protein induces IL-6 gene expression.

As mentioned, MCFG induces IL-6 expression, resulting in slightly inhibiting PRV RNA replication and significantly reducing syncytial formation, but the antibody to p17 is only significantly inhibiting syncytial formation. These suggest that IL-6 likely directly interferes with syncytial formation and has a minimal impact on PRV RNA replication, both of which are fundamental PRV processes associated with IL-6 expression. The difference between the effects of MCFG and anti-p17 antibodies is that MCFG can also inhibit other PRV proteins. In comparison, anti-p17 may be specific to p17 alone, so it does not affect other processes related to RNA replication. The inhibition of PRV RNA replication by MCFG was mediated through its interaction with other target proteins, as identified by a molecular docking study, which showed that MCFG binds to the nonstructural replication protein (sigmaNS; σNS gene) as the second target. Previous research indicated that the function of the σNS protein of *orthoreovirus* is essential for viral replication and assembly, functioning as an RNA-binding and stabilizing agent. The σNS is crucial for the formation of viral replication factories. The structural properties enable the compartmentalization of replication machinery, thereby ensuring efficient genome replication and capsid assembly [67]. Additional insights indicate that σNS stabilizes viral RNA, safeguarding it from degradation and facilitating genome synthesis during replication. This RNA stability function underscores its significance in preserving the integrity of the viral replication process [68]. Furthermore, they illustrate how σNS assembles into oligomers that associate with viral RNAs and direct them to replication organelles. This recruitment highlights the function of σNS as a principal organizer of the viral replication apparatus, facilitating the availability of viral RNAs for transcription and incorporation into new virions [69].

In summary, this study revealed that MCFG potentially targets PRV p17, effectively inhibiting viral release and syncytial formation while slightly reducing PRV replication in A549 cells. Notably, MCFG may target additional PRV proteins beyond p17, such as nonstructural replication proteins (sigmaNS gene), as identified in a molecular docking study. Given these findings, MCFG shows promise as an antiviral treatment option for controlling PRV propagation and its associated pathogenesis.

## Supplementary Information

The supplementary information available in Supplementary information 1.xlsx

## Acknowledgments

We would like to thank the National Institute of Genetics, Japan for the Supercomputer System for RNA-Seq analysis.

## Contributions

EH contributed to the study conception and design. Material preparation, data collection, and analysis were performed by WB, AI, and EH. The first draft of the manuscript was written by WB, and all authors commented on previous versions of the manuscript. All authors read and approved the final manuscript.

## Data Availability

Data is included in the manuscript or supplementary information.

## Declarations

### Ethics approval

The Ethics Review Board for Animal Experiments was approved at Nagoya University (Date 2024/10/04; approval number: A240118-004).

